# Does strain-level persistence of lactobacilli in long term back-slopped sourdoughs inform on domestication of food-fermenting lactic acid bacteria?

**DOI:** 10.1101/2024.09.26.615182

**Authors:** Vi D. Pham, Zhaohui S. Xu, David J. Simpson, Justina S. Zhang, Michael G. Gänzle

**Affiliations:** University of Alberta, Dept. of Agricultural, Food and Nutritional Science, Edmonton, Canada

**Keywords:** Fermentation control, lactic acid bacteria, *Lactobacillus*, whole genome sequencing, single nucleotide polymorphisms, rate of mutation, speciation, domestication

## Abstract

Sourdoughs are maintained by back-slopping over long time periods. To determine strain-level persistence of bacteria, we characterized 4 sourdoughs from three bakeries over a period of 3.3, 11, 18 and 19 years. One sourdough included isolates of *Levilactobacillus* spp. and *Fructilactobacillus* spp. that differed by fewer than 10 SNPs from the isolates obtained 3.3 years earlier and thus likely represent the same strain. Isolates of *Lv. parabrevis* differed by 200 – 300 SNPs, their genomes were under positive selection, indicating transmission from an external source. In two other sourdoughs, isolates of *Fl. sanfranciscensis* that were obtained 11 and 18 years apart differed by 19 and 29 SNPs, respectively, again indicating repeated isolation of the same strain. The isolate of *Fl. sanfranciscensis* from the fourth sourdough differed by 45 SNPs from the isolate obtained 19 years prior. We thus identified strain-level persistence in 3 out of 4 long-term back-slopped sourdoughs, making it possible that strains persisted over periods that are long enough to allow bacterial speciation and domestication.

**Importance:** The assembly of microbial communities in sourdough is shaped by dispersal and selection. Speciation and domestication of fermentation microbes in back-slopped food fermentations has been documented for food fermenting fungi including sourdough yeasts but not for bacteria, which evolve at a slower rate. Bacterial speciation in food fermentations requires strain-level persistence of fermentation microbes over hundreds or thousands of years. By documenting strain-level persistence in three out of four sourdoughs over a period of up to 18 years, we demonstrate that persistence over hundreds or thousands of years is possible. We thus not only open a new perspective on fermentation control in bakeries but also support the possibility that all humans, despite their cultural diversity, share the same fermentation microbes.

## Introduction

Sourdough is used as a leavening agent or baking improver for bread making in households and in artisanal and industrial bakeries (1). Sourdoughs that are maintained for baking are almost invariably back-slopped, i.e. the fermentation is initiated by using a portion of the previous batch as an inoculum. The practice of back-slopping was first recorded about 2000 years ago (2) but is likely much older for a simple reason: back-slopped sourdoughs are suitable for use as leavening agents but spontaneous sourdoughs are not (3). Breads other than flatbreads thus required the use of back-slopped sourdoughs until brewer’s yeast or baker’s yeast became available as leavening agent. Continuous back-slopping for periods of more than 100 years is documented for several sourdoughs (4). Back-slopping of communities of fermentation microbes exerts tight control on the composition of microbial communities, which converge at the genus- or even at the species level (3, 5, 6). Globally, virtually all bakeries that use sourdough as the sole leavening agent ferment with *Fructilactobacillus sanfranciscensis* (3, 5). Sourdoughs that are used to acidify yeast-leavened rye doughs predominantly include *Lactobacillus* and *Limosilactobacillus* species (3, 5). Improved fermentation control requires knowledge on whether these microbial communities are functionally stable, i.e. they are composed of different strains or species over time that express the same metabolic traits, or phylogenetically stable, i.e. they are composed of strains that permanently persist. Functional stability has been shown for two industrial sourdoughs that included different reutericyclin-producing strains of *Limosilactobacillus reuteri* (7) and different *Lactobacillus* species expressing the extracellular fructanase FruA (8) over periods of several years. In sourdoughs that are functionally stable, the composition of microbial communities is determined by dispersal and selection (9, 10). For example, functional stability of sourdoughs that are used for acidification of yeast-leavened rye breads is dependent on transmission of lactobacilli and limosilactobacilli. Measures that disrupt transmission paths from their establishment niche, the gut of birds and domestic animals (11, 12), likely also decrease the microbial diversity of these sourdoughs and their resilience to perturbation.

Phylogenetic stability of bacterial communities in fermented foods is supported by the observation that several bacterial taxa that dominate in back-slopped food fermentations include phylogenetic clades that exclusively consist of isolates from fermented foods. One example is *Lactococcus lactis*, which separates in two clades, one of which includes plant and dairy isolates while the other exclusively includes dairy isolates from back-slopped mesophilic starter cultures (13, 14). A second example is *Lactobacillus delbrueckii* subsp. *bulgaricus*, which is also used as dairy starter culture. All isolates of *L. delbrueckii* subsp. *bulgaricus* were obtained from thermophilic fermented dairy products (15). Long-term strain level persistence of bacteria in back-slopped food fermentations as an absolute prerequisite for domestication and speciation, however, has not been documented experimentally.

The determination of whether an isolate that was isolated from a back-slopped food fermentation is offspring of an earlier isolate from the same fermentation and thus represents the same strain requires reliable tools for strain-level identification of bacteria. The gold standard of strain level identification is whole-genome sequencing (WGS) and quantification of single-nucleotide polymorphisms (SNPs), which has widely used in investigations of outbreaks of (foodborne) disease (16, 17). Current pipelines use Illumina as the standard platform for genome sequencing (21). The long read sequencing platform Nanopore has emerged as an affordable approach to sequence genomes with an accuracy that is comparable to Illumina sequencing (18). The bioinformatic tools for SNP calling of long read sequencing, including deep learning-based SNP callers such as DeepVariant and Clair3, are rapidly developing and surpassing other SNP callers in performance (19). These advances facilitate genome sequencing and SNP calling to identify strains of food fermenting lactobacilli. It was therefore the aim of this study to analyse strain level persistence in three sourdoughs that were back-slopped in four bakeries and sampled over a period of 3.3 to 19 years. Strain level identification was achieved by genome sequencing and quantification of pairwise SNPs.

## Results

### Identification of fermentation microbes in three sourdoughs

The sourdough A from bakery A was sampled in 2019, 2022 and 2023; in addition, a rye sourdough maintained in the same bakery was sampled in 2023. The sourdough B of bakery B was sampled in 2012 and 2023. Sourdoughs A and B were maintained over the entire observation period without change of ownership or location.. The sourdough C from bakery C was sampled in 2005. In 2009, sourdough C was passed on to baker D that took over operations as well as the sourdough (sourdough C’). The bakery moved to a new facility in 2022. After 2009, baker C continued to use sourdough C to bake bread for personal use (extensive baking about once per week). This sourdough C’’ was sampled in 2024. Sourdoughs B, C, C’ and C’’ consisted of one strain of *Fl. sanfranciscensis* which accounted for more than 99% of the viable bacterial cell counts. Strains of *Levilactobacillus parabrevis* dominated both sourdough A and the rye sourdough from bakery A (Table 1). *Pediococcus parvulus* was the second or third abundant species in sourdough A and the rye sourdough maintained in bakery A. Sourdough A and the rye sourdough of bakery A differed with respect to minor members of the community. The wheat sourdough contained strains of *Levilactobacillus* and *Fructilactobacillus* which could not be assigned to known species, and acetic acid bacteria. The rye sourdough contained *Companilactobacillus crustorum*, *Levilactobacillus brevis* and *Fl. sanfranciscensis* as minor components.

**Table 1.**
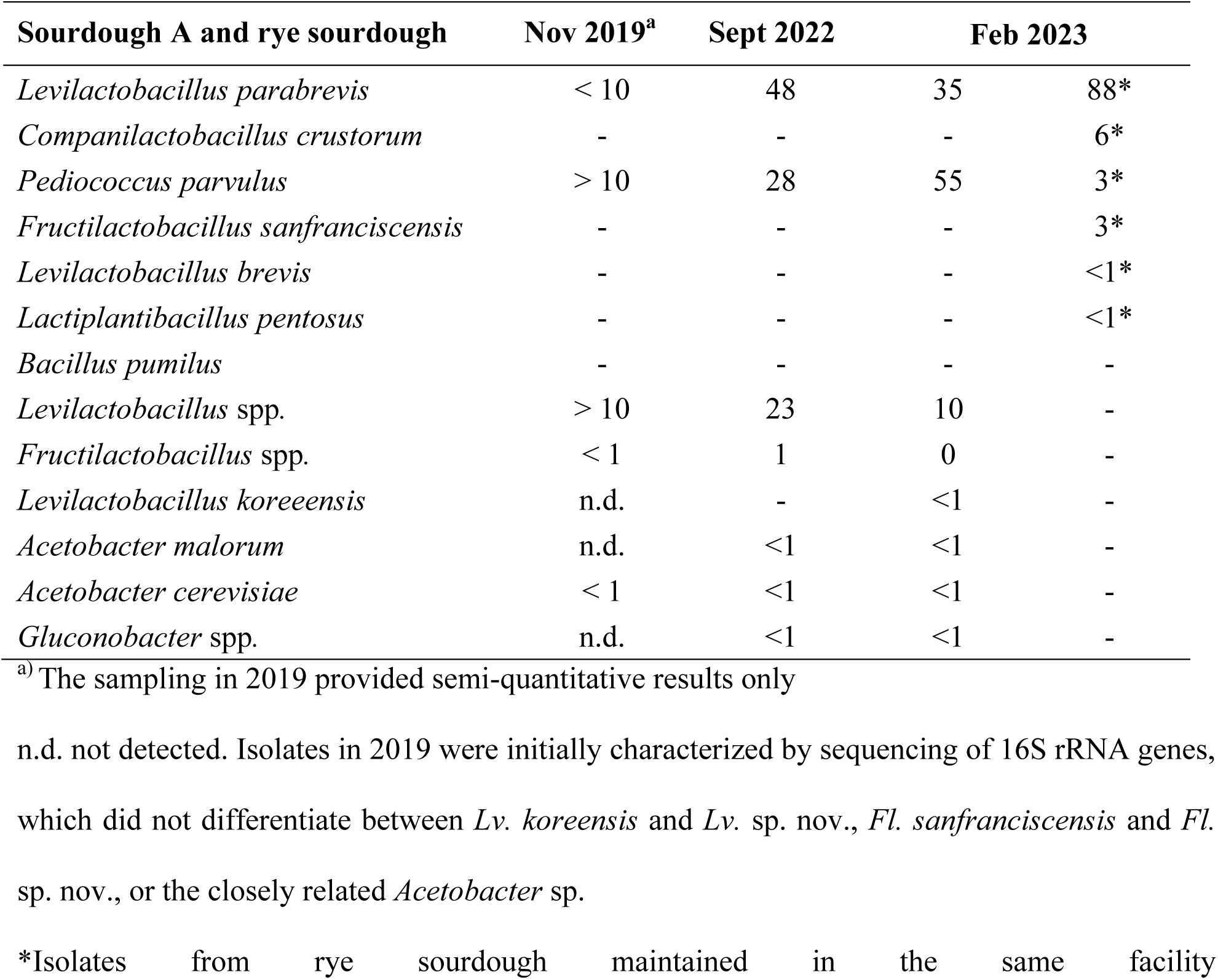
Identification of isolates by 16S rRNA Sanger sequencing. Numbers show approximate abundance (%) by colony morphology. Hyphen indicates absence and n.d. is not determined. Approximate abundance of bakery A isolates were not accessed during the Nov. 2019 sampling.

| Sourdough A and rye sourdough | Nov 2019<sup>a</sup> | Sept 2022 | Feb 2023 |  |
| --- | --- | --- | --- | --- |
| *Levilactobacillus parabrevis* | < 10 | 48 | 35 | 88* |
| *Companilactobacillus crustorum* | - | - | - | 6* |
| *Pediococcus parvulus* | > 10 | 28 | 55 | 3* |
| *Fructilactobacillus sanfranciscensis* | - | - | - | 3* |
| *Levilactobacillus brevis* | - | - | - | <1* |
| *Lactiplantibacillus pentosus* | - | - | - | <1* |
| *Bacillus pumilus* | - | - | - | - |
| *Levilactobacillus* spp. | > 10 | 23 | 10 | - |
| *Fructilactobacillus* spp. | < 1 | 1 | 0 | - |
| *Levilactobacillus koreeensis* | n.d. | - | <1 | - |
| *Acetobacter malorum* | n.d. | <1 | <1 | - |
| *Acetobacter cerevisiae* | < 1 | <1 | <1 | - |
| *Gluconobacter* spp. | n.d. | <1 | <1 | - |

<sup>a</sup>) The sampling in 2019 provided semi-quantitative results only
n.d. not detected. Isolates in 2019 were initially characterized by sequencing of 16S rRNA genes, which did not differentiate between *Lv. koreensis* and *Lv. sp. nov.*, *Fl. sanfranciscensis* and *Fl. sp. nov.*, or the closely related *Acetobacter* sp.
\*Isolates from rye sourdough maintained in the same facility

To complement culture-based analyses on the composition of the microbial community, 16S rRNA genes were amplified from DNA isolated from sourdoughs and sequenced. The sequencing result qualitatively matched the culture-based identification but with a distorted relative abundance of several taxa (Fig. S1). In particular, *Fructilactobacillus* sequences were strongly over-represented in the rye sourdough of bakery A.

### SNP comparison between Nanopore-and Illumina-sequenced genomes

To confirm the accuracy of the workflow that was developed in this study for SNP calls in Nanopore-sequenced genomes, we analysed the number of SNPs between Nanopore reads (duplex reads with Q score ≥ 20 basecalled with super accurate model) and Illumina-sequenced genomes across different read depths for 3 organisms (Fig. 1). The Illumina genomes were sequenced with a coverage of more than 100-fold. The Nanopore reads were randomly sampled into a specific coverage to avoid sampling bias. Nanopore-sequenced genomes of *Weissella cibaria* FUA3120 and *Listeria monocytogenes* FSL C1-056 differed in 0 or 5 SNPs from the Illumina genomes when the sequencing depth was 20-fold coverage or higher. For *Lactiplantibacillus plantarum* TMW1.460, an 80-fold or higher coverage was required to consistently obtain a difference of 2 SNPs (Fig. 1). The fluctuation between 2 and 9 SNPs in the *Lp. plantarum* TMW1.460 resulted from random sampling of the reads. Overall, this validates the workflow for SNP calling with fewer than 5 false-positive SNPs per genome.

**Figure 1.**
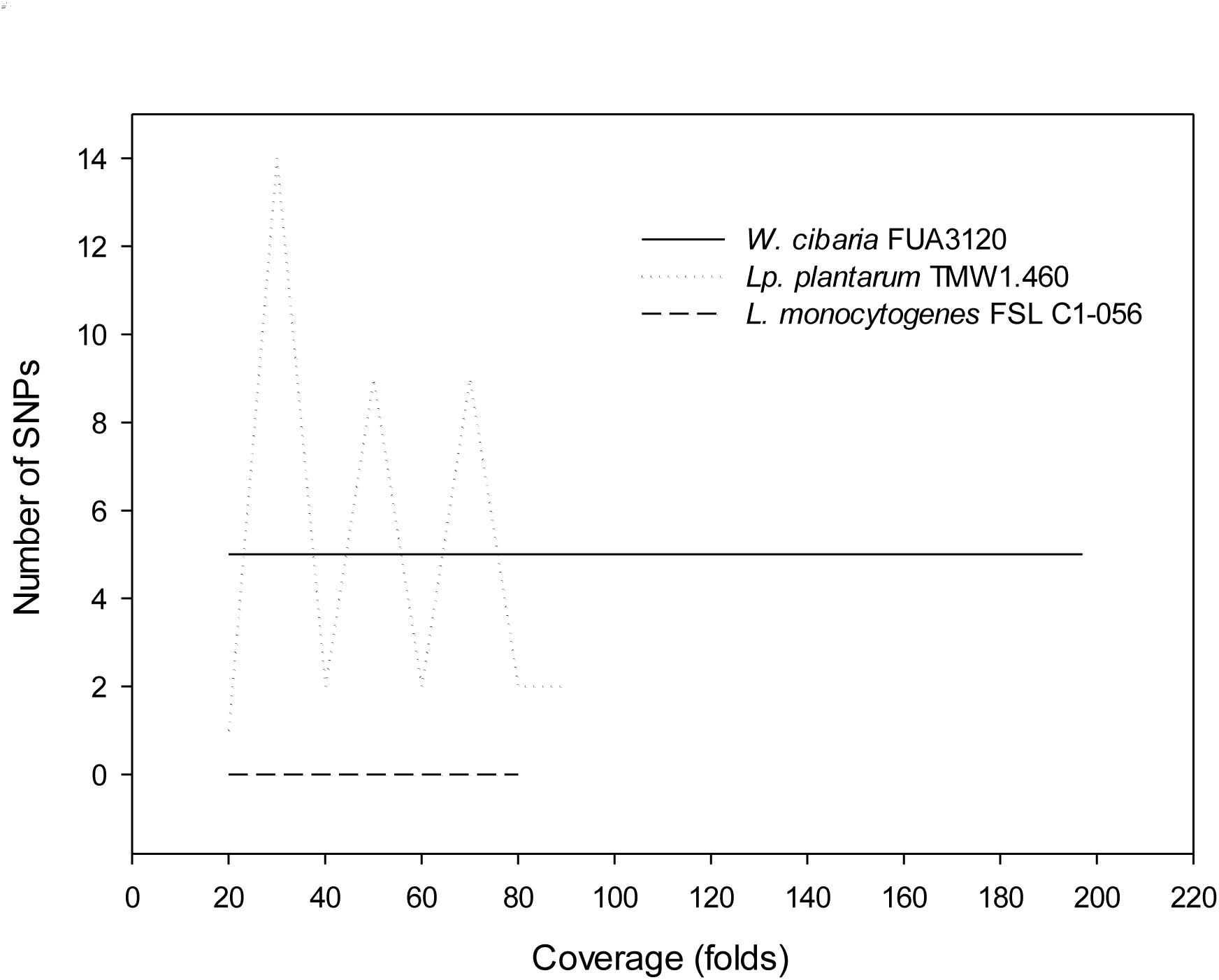
The discrepancy between Illumina and Nanopore sequencing across coverages in three food associated bacterial strains isolated independently. Reads were randomly subsampled into specific coverages. Nanopore reads of the same strain were mapped against its Illumina genome. SNPs were called by Clair3 and visualized on IGV for potential false positive SNPs.

### SNP analysis of sourdough isolates obtained from four sourdoughs over a period of 3 to 19 years

Isolates obtained at each sampling time were selected for SNP analysis (Table S1). For SNP analysis, the earliest sampled isolate in each sourdough obtained in 2019 and 2012 was used as a reference. For sourdough C, the 2023 isolate from sourdough C’ was used as reference because the genome sequence of the 2005 isolate was available only as metagenomic assembled bin. Revival of the 2005 isolate in sourdough did not result in any viable cell of *Fl. sanfranciscensis* determined by colony PCR and species-specific primers. However, DNA from *Fl. sanfranciscensis* was present in the initial sourdough containing the frozen stock, water and flour. Metagenomic reads resulting from sequencing of sourdough DNA and taxonomically assigned to *Fl. sanfranciscensis* were mapped against the 2023 genome for SNP calling. The reference genomes were assembled and polished with minimum 135-fold coverage. The mean read depth aligning to the reference genomes was 210-fold across all isolates.

Table 2 shows the SNPs of each isolate relative to its reference indicated by the year of sampling. The SNP distance matrix for isolates of each sourdough is reported in Tables S2-S5. The number of SNPs varies among species. In sourdough of bakery A, isolates of *Levilactobacillus* spp. and *Fructilactobacillus* spp. from 2023 differed in fewer than 10 SNPs from the respective isolates obtained in 2019. The 2023 isolates of *Pc. parvulus* differed by 30 SNPs from those isolates obtained in 2022 and in 2019. The highest number of SNPs were observed in isolates of *Lv. parabrevis*, ranging from 189 to 321. Using one 2022 isolate as a reference to an isolate obtained in 2023 identified 313 SNPs (Table 2, Table S2). In the sourdoughs B, C and C’, the respective *Fl. sanfranciscensis* isolates differed by 19 and 29 SNPs. The 29 SNPs were observed with a reference genome that was of relatively low quality. The culture stock from 2005 could no longer be resuscitated on agar media. After an unsuccessful attempt to recover the strain after a sub-culture in wheat sourdough, metagenomic sequencing of the sourdough resulted in ∼6.1 Gbases, of which ∼3.3 Mbases were classified as *Fl. sanfranciscensis*. Mapping of the metagenomic reads resulted in 2.5-fold read depth and covered 80% of the *Fl. sanfranciscensis* genome. The isolate from sourdough C’’ differed from the isolate from sourdough C’ (30 years of separate evolution) by 89 SNPs and from the isolate from sourdough C (19 years of evolution) by 45 SNPs.

**Table 2.** SNP distance between each recently sampled strain relative to the first sampled strain (indicated by sampling time and sourdough).

| Sourdough |  | Isolate indicated by year (sourdough) |  |  |
| --- | --- | --- | --- | --- |
|  |  | 2019 | 2022 | 2023 |
| A | *Lv. parabrevis* |  | 189, 321 | 319 (313)<sup>a</sup>, 318, 205*, 210* |
|  | *Levilactobacillus* spp. |  | 3 | 9 |
|  | *Fructilactobacillus* spp. |  | 6, 8 | - |
|  | *Pc. parvulus* |  | 30 | 30, 30* (14) |
| B |  | 2012 |  | 2023 |
|  | *Fl. sanfranciscensis* |  |  | 19 |
| C, C' and C'' |  | 2005<sup>b</sup> | 2024 (C''), 2023 (C') |  |
|  | *Fl. sanfranciscensis* |  | 45, 29 |  |
|  |  | 2023 (C') | 2024 (C'') |  |
|  | *Fl. sanfranciscensis* |  | 89 |  |

\* An asterisk designates isolates from the rye sourdough; all other isolates are from the wheat sourdough that is maintained separately in the same facility.
<sup>a)</sup> Numbers in brackets compare the strains to strains isolated in 2022.
<sup>b)</sup> The culture stock of the 2005 isolate in the FUA strain collection was no longer viable. The metagenomic reads from a sourdough started with the content of the 2 mL freezer vial were classified with 0% contamination and 80% coverage to the reference genome.

### Signatures of adaptation of genomes of *Lv. parabrevis* isolates

To determine whether the high number of SNPs in strains of *Lv. parabrevis* was a result of a high mutation rate of the same strain or repeated introduction of the closely related environmental strains, we determined whether the genomes are under positive selection (Table 3). The estimated non-synonymous to synonymous rate ratio (dN / dS) was 1.003, demonstrating (*P* ≤0.05) that the isolates are under positive selection. At least one of the DNA repair gene (*mutL / M / S / Y*) was impacted in every genome with SNPs resulting either in premature stop codons and / or altered protein function. An exceptionally high transition to transversion (Ti / Tv) ratio of 22 - 158 and missense to silent mutation ratio of about 2 also indicate amino acid changes towards adaptation to the new environment (24). SNP annotation of the type strain of *Lv. parabrevis* and the selection pressure of this branch revealed a Ti / Tv ratio of 5 and a dN / dS ratio of 0.86, respectively, which are close to the values that are expected for genomes that experience neutral selection. The evolutionary pressure is thus specific to the isolates from sourdough A and not a general property of the species.

**Table 3.** Estimated nonsynonymous to synonymous rate ratio (dN / dS), transition to transversion ratio (Ti / Tv), and effects of SNPs in coding regions and DNA repair genes in strains of *Lv. parabrevis* isolated from sourdough of bakery A.

| Strains by year | dN / dS | Ti / Tv | Mutations on protein |  | Mutations in *mutL/M/S/Y* |  |  |
| --- | --- | --- | --- | --- | --- | --- | --- |
|  |  |  | Missense /<br>silent | Premature<br>stop codon<br>(%) | Missense | Silent | Premature<br>stop codon |
| 2022 | 1.003 | 93.5 | 1.9 | 0 | 1 | 0 | 0 |
|  |  | 106 | 2.3 | 1.1 | 1 | 0 | 1 |
| 2023 |  | 158 | 2.1 | 1.5 | 1 | 0 | 1 |
|  |  | 64.2 | 2.2 | 1 | 0 | 0 | 1 |
| 2023* |  | 28.3 | 2.4 | 0 | 0 | 1 | 0 |
|  |  | 22.2 | 2.6 | 0 | 0 | 1 | 0 |

<sup>\*</sup> An asterisk designates isolates from the rye sourdough; all other isolates are from the wheat sourdough that is maintained separately
in the same facility.

## Discussion

The sourdoughs that were analysed in this study were maintained in different fermentation schemes. Sourdoughs B, C and C’ are maintained for use as sole leavening agents in bakeries (3). Sourdough B has been back-slopped twice a day with 10 % inoculum at room temperature. Sourdough C has been back-slopped twice a day until 2009 with fermentation between 12 and 18 °C (fermentation in a cool basement) and once a day with 25 % inoculum and fermented at 15 °C (temperature control in an incubator) since 2009. As expected, sourdoughs B, C and C’ included *Fl. sanfranciscensis* as a dominant representative of the bacterial community (3, 5). *Fl. sanfranciscensis* was maintained in sourdough C’’ although the sourdough was back-slopped much less frequently to bake only about once per week for personal use at the household level. Sourdough A is very distinct from any other sourdough as it is propagated with exceptional fermentation parameters, i.e. constant fermentation below 10 °C for a fermentation time of 6 d. The rye sourdough of bakery A follows the same propagation but includes a 1 d fermentation cycle at room temperature between the fermentation cycles below 10 °C. In keeping with the exceptional propagation conditions, the microbial communities in sourdoughs of bakery A were also exceptional. Sourdough A included one strain each of *Levilactobacillus* spp. and *Fructilactobacillus* spp. that could not be assigned to a known species. The 15 species in the *Lactobacillaceae* which were described with sourdough isolates as type strains were described before 2013 (5, 20) or with isolates obtained before 2013 (21), indicating that the diversity of sourdough lactobacilli is all but fully described with isolates that are currently available. The isolation of two novel species in a single sourdough thus also reflects the unprecedented propagation conditions (3, 5).

Knowledge of the propagation conditions allows a reasonably accurate estimation of the number of bacterial generations between the first and the last sampling of each of the three sourdoughs. In 3.3 years, the strains in sourdough A grew for approximately 200 generations. The strains from sourdough B grew for approximately 28,000 generations in 11 years and the strain from sourdough C’ grew for an estimated 29,000 generations in 18 years.

### Validation of the workflow for SNP calling using Nanopore sequencing

Regulatory agencies developed workflows for SNP analyses to identify strains of pathogens using Illumina data (16, 17). As the Nanopore sequencing platform rapidly emerges, recent studies investigating the discrepancy in SNPs between Nanopore and Illumina data concluded that both platforms provide comparable results for the pathogens *Mycobacterium tuberculosis, Salmonella enterica, Escherichia coli,* and *L. monocytogenes* (22–25). These studies also performed a series of read processing and filtering steps but used SNP callers that are generic or previously developed for short read data. We used Clair3, a deep learning SNP caller designed specifically for Nanopore reads with the highest F1 score among currently available SNP tools (26, 27). We basecalled the duplex reads using the sup model and demonstrated that at 20-, 50-, and 80-fold coverage, the SNPs are 0 – 14, 0 – 9, and 0 – 5, respectively. In this basecalling model with the duplex reads, Clair3 achieved the F1 score of ≥ 99.99% for SNP calls at read depth of ≥ 25 when benchmarked against 14 bacterial species (19).

### Strain level persistence of sourdough isolates

Strain level identification of foodborne pathogens and nosocomial pathogens in outbreak investigations of foodborne disease or pathogenic transmission has been determined by a cut-off of about 21 SNPs (16, 17). The same threshold has also been used to determine persistence of *L. monocytogenes* in food processing facilities (28, 29). Since a universal definition of bacterial strains is not established, strain identification depends on the context. For example, while a SNP cut-off of 21 reliably identifies strains of *L. monocytogenes* in outbreak investigations (16), strains of the same species accumulated fewer than 10 SNPs over a period of up to 17 years of persistence in food processing facilities (28). In the context of this study, the strain level identification informs on the probability of persistence versus repeated contamination with closely related strains. Estimating the limit of detection of the number of SNPs in analogy to analytical chemistry as baseline + three standard deviations correspond to 10 SNPs that confidently conclude on differences between two genomes (Fig. 1) (30). The pairs of isolates of *Levilactobacillus* sp. nov. and *Fructilactobacillus* sp. nov. differ by fewer than 10 SNPs and are thus very likely representatives of the same strain. The isolates of *Fl. sanfranciscensis* from sourdoughs B, and C and C’ differ by 19 and 29 SNPs, respectively, and are also likely the same strain. The isolates are separated by 11 and 18 years of evolution and the latter SNP is inflated due to low quality meta-assembled genome and the low read depth. The isolate of *Fl. sanfranciscensis* from sourdough C’’ differed by 45 SNPs from the isolate from sourdough C and by 89 SNPs from the isolate from sourdough C’. The higher number of SNPs may reflect a higher rate of mutation after the sourdough transitioned from daily baking to extensive (about weekly) baking, or indicate that the sourdough C’’ is populated by a different strain. Strain level persistence of *Pc. parvulus* with 30 SNPs in 3.3 years corresponding to 200 generations is possible but not more likely than repeated contamination. Continuation in sampling and further analyses are needed to determine whether the strain persists over time, or not.

What are possible mechanisms for the strain-level persistence of *Fl. sanfranciscensis*? As outlined in the introduction section, the use of sourdough as sole leavening agent exerts control over the microbial communities in sourdough – an acceptable product that meets consumer expectations is obtained only is the metabolic activity of sourdough microbes is maximised (31). *Fl. sanfranciscensis* is not known to occur in any niche other than wheat or rye sourdoughs that are maintained for use as sole leavening agent (32), which may indicate that is has evolved to adapt to this niche and out-competes all other bacteria. The ability of *Fl. sanfranciscensis* to outcompete other microbes apparently facilitates its long-term strain-level persistence. Sourdoughs dominated by *Fl. sanfranciscensis* generally include other lactobacilli with a much lower abundance (5, 33) but our experimental design did not account for these strains.

The *Lv. parabrevis* isolates with 200 – 300 SNPs difference between isolates obtained in 2019 and 2022 or 2023, and particularly with 313 SNPs between isolates obtained in 2022 and 2023 are likely repeat contaminants (Table 2). Positive selection as indicated by the dN / dS ratio in conjunction with a remarkable mutation bias towards transition and high missense to silent mutation ratio indicate that these organisms adapt to the sourdough environment after transitioning from another environment (34, 35). Transition bias is well documented in organisms adapting to a new environment with increased rate of spontaneous mutations. For example, *E. coli* continuously accumulated mutations of which > 90% were transitions during adaptation to a 3-year stationary phase (35, 36). Once adaptation has taken place, selection favoured the population with a reduced mutation rate (37). The *Lv. parabrevis* isolates are thus likely closely related environmental strains that are repeatedly introduced to the sourdough (Table 2, Table S2).

### Strain level persistence, speciation and domestication of food fermenting bacteria

Domestication of food fermenting microbes is documented for several fungi including brewer’s yeast, sourdough yeasts, and *Penicillium roqueforti* (38–40). Eukaryotic organisms evolve at a faster rate than prokaryotes, however, and genetic and physiological signs of domestication are observed after a few hundred years, i.e. over a timeline that is documented by a historical record (38, 41). Bacteria evolve at a slower rate, i.e. over a timeline that is not documented by the historical record but requires archeological findings. Specifically, all contemporary isolates of *Fl. sanfranciscensis* are essentially clonal and would be considered to be the same strain when judged on the criterion >99.99% ANI (Fig. 2) (42). When extrapolating the mutation rate based on the SNP differences over periods of 11 or 18 years and the rate of mutation of *L. monocytogenes* over a comparable period (28), a period of about 2000 to 10,000 years or more is required for evolution of phylogenetic lineages of prokaryotes that diverge at 98% ANI or less. Current knowledge does not allow to align the history of food fermentations and the rate of bacterial evolution more precisely as an archeological record for genomes of food fermenting bacteria is lacking (Fig. 2). A first and absolute prerequisite for domestication of bacteria is thus continuous back-slopping of sourdoughs, and strain-level persistence of fermentation microbes in sourdoughs for 2000 – 10,000 years. We identified persisting strains in three out of four sourdoughs, indicating that strain level persistence is more likely than not. Strain level persistence in back-slopped food fermentations over a period of about 400 years is also supported by the domestication events in *P. roqueforti* and *Saccharomyces cerevisiae* (38–40). Sourdoughs have a documented or credible age of 100 – 200 years (4) (Fig. 2) but may be substantially older. Bakers pass their sourdoughs on to the next generation, as was the case for sourdough C. Likewise, sourdoughs that are currently used in San Francisco and in the Yukon territory can credibly be dated to the gold rushs of 1849 and 1898, respectively, but probably have their roots on the U.S. East coast and may have been brought by immigrants from Europe. Transfer and transport of sourdoughs is a necessity for baking as spontaneous sourdoughs are not suitable as leavening agents. Even though experienced bakers address dispersal limitations by inoculation with plant material or manure (43), it takes days or weeks to build an active sourdough from scratch. In short, whenever sourdough-leavened loaves of bread or steamed bread are desired, mortal bakers bake with immortal sourdoughs. Likewise, North American dairy starter cultures are thought to all originate from few starter cultures that were brought from Europe (44). When assuming that strain-level persistence is observed more often than not, i.e. occurs in many or most sourdoughs that are used as sole leavening agent, a timeframe of more than 2,000 years of continuous back-slopping is possible: this prerequisite for domestication is possibly met. Domestication is also supported by phenotypic and genotypic differences of *Fl. sanfranciscensis*. The organism has the smallest genome size of all fructilactobacilli and the highest density of rRNA operons per Mbp genome, and is distinguished from other fructilactobacilli by the ability to metabolise maltose. The small genome size, the high density of rRNA operons and the ability to use sucrose and maltose all support rapid growth in wheat sourdoughs (20, 45, 46).

**Figure 2.**
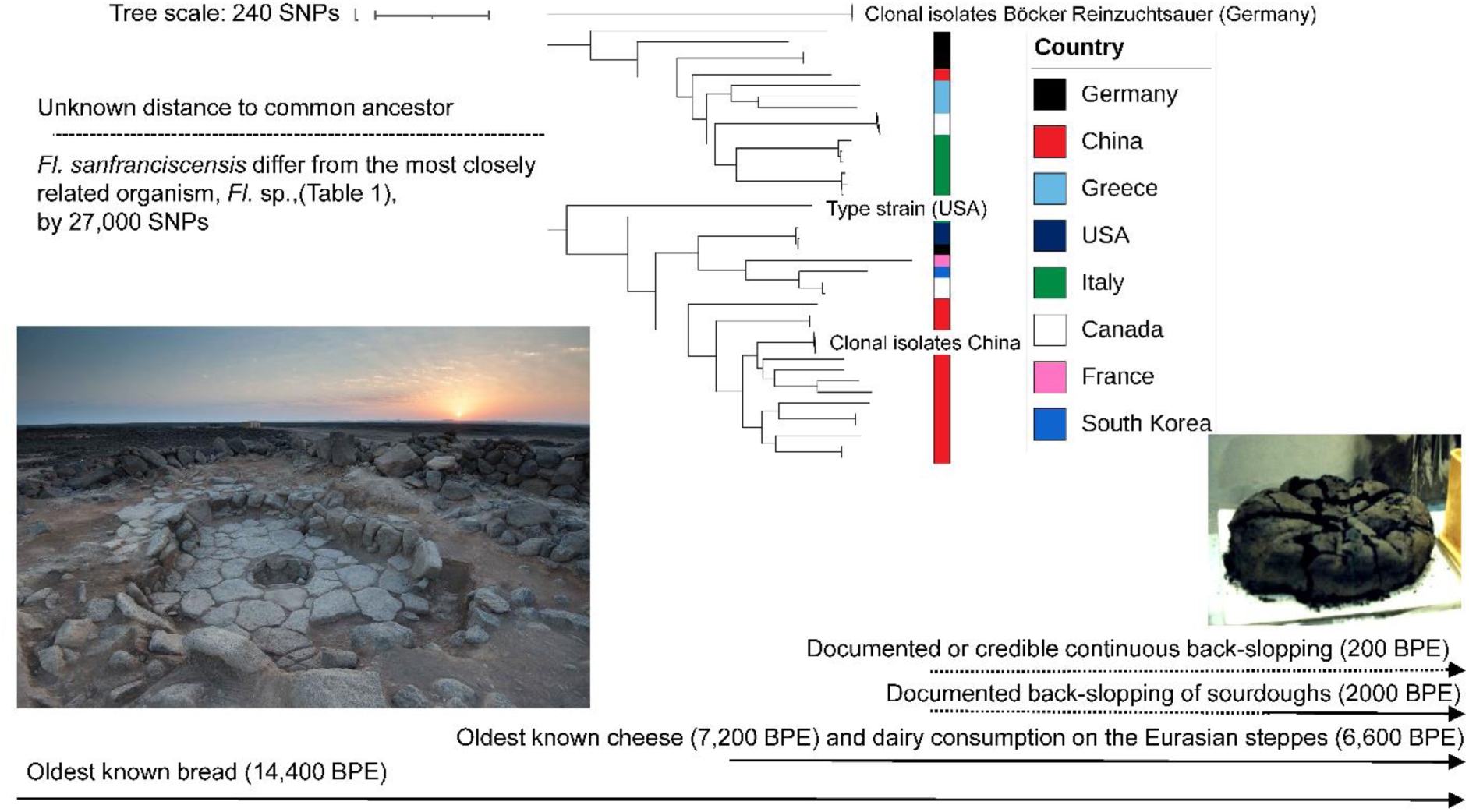
Upper panel. Unrooted SNP tree including all genomes of *Fl. sanfranciscensis* that were available on NCBI in July 2024 (Table S6). The most closely related species to *Fl. sanfranciscensis* (*Fructolactobacillus* sp., Table 1) differs by 27,000 SNPs. **Lower panel:** Documented (solid lines) and possible (dotted lines) timelines for baking (71), cheese production (72), dairy consumption on the Eurasian Steppes (48), back-slopping of sourdoughs (2) and continuous back-slopping of a sourdough (4). The picture to the lower left shows one of the fireplaces where bread-like remains were found (published under a creative commons license (71); the picture on the right shows a bread that was buried in Pompei during the eruption of the Vesuvius in 79 (picture of the authors).

A second and equally absolute prerequisite for domestication is that all isolates of the domesticated phylogenetic clade originate from a single domestication event. *P. roqueforti* and *S. cerevisiae* include several domesticated lineages that were domesticated in different locations (38, 40, 41). In the bacterial organisms for which domestication is possible or likely, i.e. *Lc. lactis*, *Lb. delbrueckii* subsp. *bulgaricus*, *Lactobacillus kefiranofaciens* subsp. *kefirofaciens* and *Fl. sanfranciscensis*, the potentially domesticated phylogenetic clades include isolates that were obtained globally (Fig. 2) (5, 14, 47). This global distribution cannot be explained by the globalization of trade and the development of starter cultures in the 20^th^ century. In addition, the presence of *Fl. sanfranciscensis* was not observed in any of the dozens of spontaneous sourdoughs for which species-level information is available (5). The topology of the phylogenetic trees is thus possible only if the organisms were distributed globally after a single domestication event about 2,000 – 10,000 years ago.

Can we identify a time in human history that made such global distribution possible? Dairying by pastoralist societies living on the Eurasian steppes 2,500 – 7,500 years ago enabled human migration both to the East and to the West (48–50). Human migration and trade across the Eurasian steppes also enabled the global distribution of goats after multiple domestication events in the Fertile Crescent, of horses after domestication on the Eurasian Steppes, and the global distribution of chicken after domestication in Thailand (51, 52). The period defined by the domestication and global distribution of goats, horses and chicken in the period of 10,000 years before present (BPE) to 3,500 BPE (51–53) is a reasonable match to the estimated time that is required for diversification of the domesticated phylogenetic clades in *Lc. lactis*, *Lb. delbrueckii* and *Fl. sanfranciscensis*. If human migration and trade supported global distribution of goats, horses and chicken, global distribution of food fermenting cultures is possible.

To add a word of caution: Bakers A and B build their sourdoughs, which both included strains of *Fl. sanfranciscensis*, from scratch. Baker B works as an instructor in a polytechnic in Edmonton, a major metropolitan area which makes transmission through physical contact with other bakers entirely possible. Baker A, however, is located in a remote and small town in the Canadian Rocky Mountains and the lack of contact to other bakers is documented by the exceptional fermentation parameters that are used for sourdough propagation. An environmental source of *Fl. sanfranciscensis* thus remains possible and conclusion on the domestication and global distribution of food microbes are dependent additional evidence, e.g. the reconstruction of genomes from archeological artefacts.

In conclusion, we established a workflow for strain-level identification by using SNP calling with Nanopore sequenced bacterial genomes. Analysis of three sourdoughs that were maintained by four bakers in as many bakeries documented strain level persistence of major representatives of the bacterial community in all three sourdoughs. The documentation of strain level persistence of major fermentation microbes in back-slopped sourdoughs informs fermentation control in sourdoughs that are maintained in bakeries. Long-term strain level persistence of lactobacilli in back-slopped food fermentations also demonstrates that one of two prerequisites for domestication and global distribution of fermentation microbes is met. Although we still lack critical evidence that some food fermenting microbes are indeed offspring of cultures that were domesticated, and distributed globally, after single domestication events about 2,000 to 10,000 years ago: The thought that humankind in all of its cultural and ethnical diversity shares the same food cultures is comforting.

## Materials and methods

### Collection of sourdoughs

Sourdoughs A and C’’ were shipped in a tight container from the original location at ambient temperature and examined for experiment within 7 d from the shipping date. Sourdoughs B, C and C’ were collected into a sterile sealable tube using a sterile wooden tongue at the original location, stored at ambient temperature and assessed within 3 h. Unless otherwise noted, all sourdough were maintained with wheat flour.

### Isolation of lactic acid bacteria from sourdoughs

Serial dilutions of sourdough homogenized in 1% peptone and 0.9% NaCl were plated on modified MRS6 (mMRS6) (54), mMRS5 (55), and glucose-M17 (55) agar supplemented with 100 mg / L of cycloheximide to inhibit growth of yeasts. While no single medium recovers all microbes from sourdough fermentations, the use of several suitable media was shown to isolate all microbes (55) The mMRS and M17 plates were incubated anaerobically and aerobically, respectively, at 30 °C for 48 h. Representative colonies of each morphology (square root of total number of the colonies on the countable plate) were re-streaked on to a corresponding agar for at least 3 times for purification. Culture of purified isolates was stored in 30% glycerol at -80 °C.

### Revival of 2005 isolate from bakery C

Since streaking and inoculation of the frozen stock onto mMRS6 agar and broth did not recover viable cultures – freezer failures and the need to defrost freezers exposed the culture stock led to temperature fluctuations that likely inactivated the strain - - we attempted to revive the isolate in flour and water, the best known growth medium for *Fl. sanfranciscensis*. The entire content of the frozen stock was mixed with 10 g of flour and 10 mL of sterile water. The mixture was incubated at 30 °C and backslopped every 12 h for 10 cycles. The backslopping was performed by mixing 1 g of the sourdough with 9 g of fresh flour and 6 mL of water. At 0 h and every 12 h, sourdough was also collected for cell count and DNA isolation. Sourdough DNA was extracted using the Qiagen DNeasy Blood and Tissue Kit (Qiagen, Hilden, Germany), and subjected to *Fl. sanfranciscensis* screening by PCR using the *Fl. sanfranciscensis* specific primers (5’- GGAGGAAAACTCATGAGTGTTAAG-3’, 5’- CAAAGTCAAGAAGTTATCCATAAACAC-3’) as described in (56). The colonies on the plates were also subjected to PCR screening of *Fl. sanfranciscensis* using the same primers. Sourdough DNA that was positive for *Fl. sanfranciscensis* was stored at -20 °C for metagenomic sequencing as described below.

### Identification and sequencing of isolates and sourdough microbial community

High molecular weight genomic DNA of purified isolates was extracted using the Promega Wizard Genomic DNA Purification Kit (Promega, Madison, WI, USA) and subjected to random amplified polymorphic DNA (RAPD) PCR (57) to eliminate clonal isolates. Isolates with unique RAPD profile were selected for 16S rRNA Sanger sequencing using the universal bacterial primers 27F (5′-AGAGTTTGATCCTGGCTCAG-3’) and 1492R (5′-TACGGYTACCTTGTTACGACTT-3′). Isolates with different RAPD profiles in one sampling time were selected for whole genome sequencing. Sequencing procedure remained identical for whole genome of isolates and metagenomic sequencing of 2005 isolate. Briefly, DNA quality and quantity were monitored using the NanoDrop spectrophotometer (Thermo Fisher Scientific) and Qubit fluorometer (Invitrogen). Sequencing library was prepared using the Native Barcoding Kit 24 V14 (SQK-NBD114.24) and sequenced on the Nanopore MinION Mk1B platform on the R10.4.1 flow cells.

For characterization of the sourdough microbial community by 16S rRNA amplicon sequencing, DNA was extracted from sourdough using the Qiagen DNeasy Blood and Tissue Kit (Qiagen, Hilden, Germany). DNA quality and quantity were monitored as described and sequenced using Nanopore MinION Mk1B platform on the Flongle R9.4.1 flow cells. Sequencing library was prepared using the 16S Barcoding Kit 24 (SQK-16S024).

### Genomic and metagenomic analysis

For genomic analyses of isolates, sequencing reads were basecalled using Dorado duplex v0.5.3 model dna_r10.4.1_e8.2_400bps_sup@v4.3.0. Basecalled reads with Q score ≤ 20 and length ≤ 500 bp were eliminated using NanoPack2 chopper v0.7.0(58) Adapters were trimmed from the filtered reads using Porechop_ABI v0.5.0 (59). Quality of the reads were monitored using FastQC v0.12.1 and NanoPlot v1.42.0. For genome assembly, high quality trimmed reads were *de-novo* assembled using Flye v2.9.3 (60) and polished using Medaka v1.11.3.

Metagenomic reads of the 2005 isolate were processed as described above and taxonomically classified using BugSeq with the NCBI nt database (61). Reads assigned to species *Fl. sanfranciscensis* were extracted using a local Python script and used for SNP calling.

For microbial community characterization by 16S rRNA gene sequencing, reads were basecalled using Guppy v6.5.7 model dna_r9.4.1_450bps_hac and classified using the EPI2ME labs metagenomic workflow v2.4.1 developed by Nanopore using minimap2 classifier and the ncbi_16s_18s_28s_ITS database.

### SNP calling

To compare Nanopore and Illumina genomes, selected isolates were also sequenced on Illumina HiSeq2500 platform. Reads from both platforms were processed and quality controlled as described. Illumina reads were *de-novo* assembled using SPAdes v3.15.5 (62). High quality trimmed Nanopore reads were randomly subsampled into various coverages using Rasusa v0.8.0.(63) Reads at each specific coverage were mapped against the Illumina assembled genome where SNPs were called and filtered using Clair3 as described below.

For isolates, SNPs were called using Clair3 v1.0.5 model r1041_e82_400bps_sup_v430 supplied by Nanopore at Rerio (https://github.com/nanoporetech/rerio) and parameters set for haploid variant as recommended (--haploid_precise --no_phasing_for_fa --include_all_ctgs -- call_snp_only) (64). Briefly, high quality reads were aligned to a reference genome (isolates with the earliest sampling date) using minimap2 v2.26 with parameters -ax map-ont --secondary=no (65). Reads coverage was assessed using mosdepth v0.3.6 (66). SNP calls and read alignment were visualized on IGV (63). SNPs were filtered manually based on the criteria as described (22). The SNP distance matrix was generated using the filtered vcf files and the CFSAN SNP pipeline (67).

### Annotation of SNPs

SnpEff v5.2c was used to annotate the vcf files containing filtered SNPs (68). The effects of SNPs were classified by number into impact on coding or non-coding regions and whether the impact alters amino acid sequence (missense, silent, nonsense mutation). Each SNP was assigned into either transitions (A/T ⟺ G/C) or transversions (A/G ⟺ C/T) and the total count was used to calculate the transition to transversion (Ti/Tv) ratio.

### Analysis of selection pressure and estimation of dN / dS

The analysis consisted of 12 available genomes of *Lv. parabrevis* (5 from NCBI and 7 from this study). A total of 1981 single-copy orthologues were inferred from OrthoFinder v2.5.5 and each orthologue alignment along with its corresponding nucleotide sequence was aligned into codon using PAL2NAL (69). The codon alignment of the single-copy orthologues was analyzed for positive selection using the branch-site unrestricted statistical test for episodic diversification implemented in HyPhy v2.5.61 (70). The method tested whether positive selection has occurred at some time at any site on any branch of isolates from sourdough A (Fig. S2) where the branch- group non-synonymous / synonymous rate ratio (dN / dS) was inferred. The method reported the result of the likelihood ratio test statistic where p-value ≤ 0.05 indicated evidence of positive selection.

## Data availability

Genome sequences of isolates that were characterized in this study were deposited to NCBI with Genbank accessions numbers documented in Table S1

## Acknowledgements

The authors sincerely thank Silvio Lettrari from Kaslo Sourdough Inc. (Kaslo, BC, Canada), Alan Dumonceaux (Northern Alberta Institute of Technology, Edmonton, AB, Canada), Yvan Chartrand (Bonjour Bakery, Edmonton, AB, Canada) and Nancy Rubuliak (previously Tree Stone Bakery, Edmonton, AB, Canada) for providing the sourdoughs for analysis, for sharing the story of their sourdoughs and their enthusiasm and knowledge on sourdough baking. Rudi Vogel, Professor Emeritus at the Technische Universität München, is acknowledged for his contributions to the comparative genomics of *Fl. sanfranciscensis*, which form an important basis of this work.

Kaslo Sourdough Inc., Mitacs, and the Canada Research Chairs Program are acknowledged for funding.

